# A reformulation of the selection ratio shed light on resource selection functions and leads to a unified framework for habitat selection studies

**DOI:** 10.1101/565838

**Authors:** Simon Chamaillé-Jammes

## Abstract

1. The selection ratio (SR), i.e. the ratio of proportional use of a habitat over proportional availability of this habitat, has for long been the standard metric of habitat selection analyses. It is easy to compute and directly estimates disproportionate use. Its apparent restriction to habitat selection analyses using categorical predictors led to the development of the resource selection functions (RSF) approach, which has now become the norm.
2. The RSF approach has however led to debates and confusion. For instance, what functional form can be used remains debated, and the concept of relative probability of selection is often misunderstood.
3. I propose a reformulation of the SR demonstrating that it can be estimated in a regression context, and thus even with continuous predictors. This reformulation suggests that RSF can be seen as an intermediate step in the calculation of SR. This reformulation also clarifies some longstanding debates about RSF and data-selection/fitting practices.
4. I further suggest that SR estimates the strength of habitat selection, but that the contribution of selection in determining use, which should be more directly linked to fitness than selection per se, should be estimated by another metric, the selection effect on use (SE). SE could be estimated simply as the *difference* between proportional use and proportional availability, and can be computed from SR and a density estimation of availability.
5. I conduct a habitat selection analysis of plains zebras to demonstrate the added-value of going beyond RSF scores and using SR estimated in a regression context, and of combining SR and SE.
6. Overall, I highlight the inter-relation between various metrics used to study habitat selection (i.e., SR, other selection indices, RSF scores, marginality). I conclude by proposing that SR and SE can be the unifying metrics of habitat selection, as together they offer a comprehensive view on the strength of habitat selection and its effect on habitat use.

## Introduction

Understanding why organisms use their environment the way they do is a key aim for ecologists (Morris, 2003). Decades of habitat selection studies have aimed at testing, quantifying, and explaining the disproportionate use of some habitats by individuals. ‘Disproportionate’ is usually formally defined in comparison with the expected use of habitats by individuals that would use the available habitats at random (Manly et al., 2002). The standard approach to estimate whether habitat use is disproportionate has for long relied on selection indices. Many selection indices have been proposed (Lechowicz, 1982; Manly et al., 2002), but the most simple and commonly used is the forage ratio first proposed by (Savage, 1931, cited in Manly et al., 2002), often (and hereafter) simply referred to as selection ratio (SR). The SR for a habitat i is defined as follow:

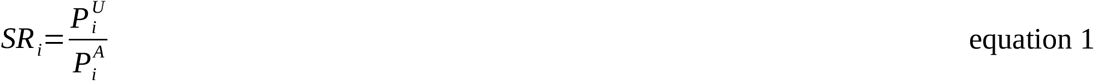

with 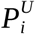 the proportion of habitat i in the set of used locations, and 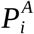 the proportion of habitat i in the set of available locations when availability is estimated using random locations. A SR_i_ value of 1 indicates that use of habitat i is proportional to its availability in the accessible landscape, and therefore a lack of selection for this habitat. A SR_i_ > 1 indicates selection, whereas a SR_i_ < 1 indicates avoidance. The common and persisting use of SR (e.g Aho & Bowyer, 2015; Basille & Fortin, 2015) shows that the SR is an index widely accepted by ecologists, probably thanks to its ease of interpretation and the clear match between its calculation and the definition of selection as a disproportionate use compared to random use.

The SR approach however appeared limited to situations in which the habitat could be defined by a limited number of categorical predictors, as one needs to be able to compute proportions over habitat categories (e.g., open vs. closed habitats). In 1990, McDonald et al. proposed a new statistical approach – resource selection functions (RSF) – to analyze the selection of habitats (or resources) defined by several variables, categorical or continuous, and their interactions (Mcdonald, Manly & Raley, 1990). This flexibility, associated with the increasing ability to integrate many habitat-defining variables thanks to advances in remote sensing, has made RSF the model of choice to estimate the strength of habitat selection.

A RSF is a function, often termed w, that transforms the density distribution of habitat i in the set of available locations into the density distribution of the same habitat in the set of used locations (Mcdonald et al., 1990; Johnson et al. 2006; Lele & Keim, 2006; McDonald, 2013), such that:

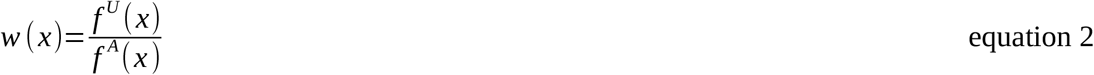

with *w*(*x*) the RSF, *f^U^*(*x*) and *f^A^* (*x*) the density distributions of used and available locations respectively. For a set of predictors x_1_ to x_n_,

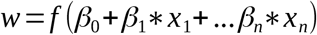

Interpreted differently, the RSF describes how the relative probability of selection of habitats varies with their characteristics defined by a set of predictors (Manly et al., 2002). As the link between eq. 1 and 2 makes clear, and as stated by (Lele, 2009), the RSF “is simply an extension of the idea of selection index to the case of continuous covariates”.

Despite its apparent simplicity, the RSF methodology has led to debates and confusion, which I now summarize:

1. RSF are commonly assumed to have an exponential form, such that

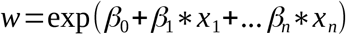 There is however little theoretical justification for the use of this specific form. The exponential RSF is the unique model when both the used and available distribution follow a normal distribution (Lele et al., 2013; McDonald, 2013), but these conditions are rarely, if ever, met. The use of exponential RSF is actually based on practicality: if one assumes an exponential RSF, the likelihood of the RSF is proportional to the likelihood of a logistic regression, with used locations being coded as 1 and available locations as 0 (Johnson et al., 2006; McDonald, 2013). Therefore, the parameters of the exponential RSF are equivalent to those returned by a logistic regression, a procedure available in virtually all statistical software. McDonald (2013) however suggested that a RSF could be any unbounded, monotonically increasing and positive function, as the only constraint he identified was that the RSF, being a ratio, should return values between [0, +Inf[. He also emphasized that Logistic, Complementary log-log or Probit functions are inappropriate because they don’t map values in the [0, +Inf[range. On the contrary, Lele et al. (Lele & Keim, 2006; Lele et al., 2013) argued however that other models, such as probit or complementary log-log functions, could actually be used, and should sometimes be favoured, to fit RSF. Thus, although most studies assume an exponential form for the RSF, it is still unclear whether other functions could be used, and how they would perform compared to exponential RSF.
2. RSF do not estimate *absolute* probability of selection (i.e. the probability of selecting an habitat when encountered (Lele et al., 2013). In RSF models, the intercept, which would allow normalizing the RSF scores to obtain probabilities of selection (Avgar et al. 2017), is considered uninterpretable because it depends heavily on how the ‘used’ and ‘available’ locations are sampled (Manly et al., 2002; Lele & Keim, 2006; McDonald, 2013). RSF are therefore only valid to “estimate *relative* probability of selecting one habitat unit with a particular set of characteristics relative to another unit with different characteristics” (McDonald, 2013). In the current RSF framework, assuming that i is the reference habitat, one can conclude that habitat j has (for instance) a higher probability of being selected than reference habitat i, but one cannot conclude about the extent to which habitats i and j are selected or avoided, compared to expectations under random use of the available habitats. Nevertheless, results from RSF analyses are often wrongly interpreted as ‘habitat j is selected’, overlooking that estimates of the strength of habitat selection are only relative. If the study of the *relative* probability of selection between habitats can in itself be sometimes of interest, often one would want to estimate whether habitats are selected, used proportionally to their availability, or avoided, irrespectively of others, in accordance with the usual definition of habitat selection being used more than expected under random use. This is most clearly seen if considering a RSF analysis with a continuous explanatory variable only, for instance distance to nearest road. The RSF will inform on whether selection increases or decreases as distance increases, but will not allow concluding about the distance at which selection shifts from attraction to random use or avoidance. It suggests that RSF cannot fulfill the original goal of most habitat selection studies. Moreover, and as emphasized by Lele et al. (2013), the interpretation of relative probability of selection is difficult: a doubling in the *relative* probability of selection does not have the same biological implication if the *absolute* probability of selection of the reference habitat is 0.004 or is 0.4. It has been claimed that *absolute* probability of selection can be estimated using resource selection probability functions (RSPF)(Lele, 2009), but not all authors agree (McDonald, 2013), and they are limited to specific situations (e.g., for instance models should contain at least one continuous predictor (Lele & Keim, 2006; Lele, 2009). Generally, very few studies have fitted RSPF models, suggesting that practitioners do not feel at ease with RSPF models.
3. RS(P)F have led to misinterpretations. As emphasized by Lele et al. (2013) who clarified it, the concept of probability of selection which underlies the RS(P)F framework is commonly misunderstood, often confounded with probability of use or probability of choice (Lele et al., 2013). The use of the logistic regression machinery, classically used in statistics to model probabilities, but in RSF analyses used to estimate the parameters of the exponential RSF and the relative probability of selection, has also created some confusion (Lele et al., 2013). It is not uncommon to see authors reporting RSF scores as if they were log-ratio of the probabilities of selection, although they are not (Lele et al., 2013; Avgar et al., 2017). Additionally, the common practice of reporting or plotting the raw coefficients of logistic regression to visualize the effect sizes is misleading because the exponentiation required to calculate the RSF scores affects the scaling.

Overall, RS(P)F are powerful tools to study complex habitat selection models, but they also have drawbacks and can be misunderstood. I re-emphasize here that the RS(P)F framework emerged from the need to integrate continuous variables in habitat selection analyses previously conducted using SR. With this in mind, I propose a new formulation of SR, mathematically equivalent to eq. 1, showing that SR are directly estimable in a classical regression framework integrating both categorical and continuous variables. I therefore argue that SR can be the standard metrics for the study of the strength of habitat selection. I complement this approach with an additional step allowing to differentiate between the strength of habitat selection and its effect on habitat use, which is mediated by habitat availability and should be more directly related to individual fitness than the strength of habitat selection (Gaillard et al., 2010).

### A reformulation of the selection ratio

The SR of any habitat i (SR_i_), when estimated using samples of use and available locations, can be reformulated as follow:

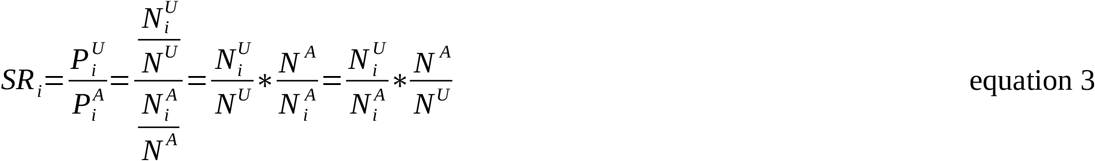

Thus, SR_i_ equals the ratio, calculated using locations falling in habitat i only, of the number of used locations 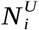 over the number of available locations 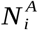, multiplied by the ratio, calculated using all locations, of the number of available locations *N^A^* over the number of used locations *N^U^*.

The ratio 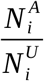 is the ratio of the numbers of locations that could be of two mutually exclusive types: be a ‘used’ location or be an ‘available’ location. Therefore, it is simply the odd of the probability that a data location falling in habitat i belongs to the set of ‘used’ locations. This is demonstrated here:

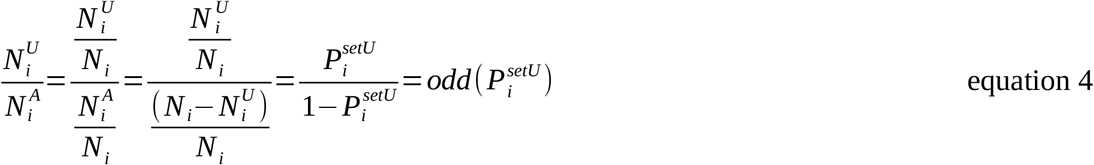

We obtain the reformulation of SR by combining eq. 3 and eq. 4:

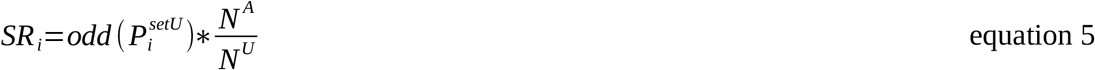

For a given analysis, *N^U^* is the total number of used locations to be analyzed, *N^A^* is selected by the analyst, and thus 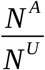 is a known scalar. Therefore, one only needs to model how oddi varies with the predictors to model how those affect habitat selection, measured using SR.

Eq. 5 provides a new justification for the use of exponential RSF estimated by logistic regression in habitat selection studies, as logistic regression is the most common statistical approach to model probabilities and odds. Logistic regression uses a logit link function to linearize probabilities (Fox, 2015), and fits:

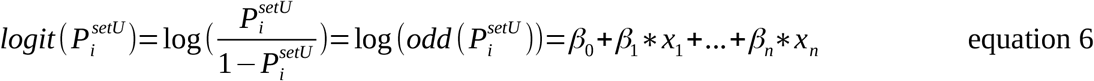

It is clear that the score 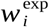 of an exponential RSF, such that 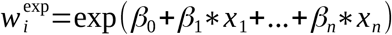, estimates 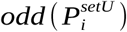, and eq. 5 shows that SR_i_ can be recovered simply:

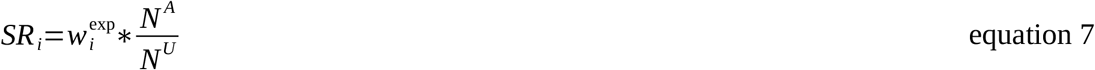

Alternatively, the RSF score can be made to be directly equal to the SR if one fits an exponential RSF using logistic regression incorporating 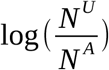 as an offset (note that the offset in logistic regression is on the logit link). Using a number of random locations equals to the number of used locations, so that 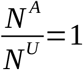, could appear an easy way to directly obtain SR without any calculations beyond RSF fitting. In most cases however, this would lead to use a relatively low number of available locations and a poor estimation of the availability of habitats (Northrup et al., 2013), and is therefore not advised. Note that, when 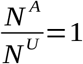 and thus w_i_ equals SR_i_, the practice of normalizing RSF scores over all available habitats leads one to obtain values that are equivalent to Chesson’s α selection index (Chesson, 1978; Lechowicz, 1982; Manly et al., 2002).

Overall, I suggest that RSF can be viewed as an intermediate step towards the calculation of SR, which could remain the central metrics to estimate the strength of habitat selection, even when one considers continuous predictors.

### Implications of the reformulation of the SR, and benefits of its use

The use of SR to estimate the strength of habitat selection naturally solves some of the aforementioned issues and debates associated with RSF modelling, and additionally shed light on another issue that had until now been overlooked when fitting RSF as hierarchical (i.e., mixed) models.

First and foremost, SR are easily interpreted and directly match the accepted definition of selection, i.e., the disproportionate use of some habitats relative to a null model of random use of the available landscape. The concept of SR may thus be viewed as conceptually more appealing than the concept of (relative) probability of selection, which is not always understood (Lele et al., 2013). The use of SR, which allow calculating selection strength for each habitat, should also prevents misinterpretation arising from the use of RSF models, which estimate differences in the strength of selection between habitats, with the strength of selection for the reference habitat remaining unknown. Note that the use of SR does not prevent the comparison of selection strength *between* habitats when these are of interest. All of this suggests that SR could be a unifying metric of the strength of habitat selection.

Second, the reformulation of SR highlights that the use of logistic regression to estimate the parameters of RSF can be viewed not as a ‘statistical trick’ based on the equivalence between the likelihoods of the logistic regression and of the use-availability or point-process models (e.g. (Johnson et al., 2006; Aarts, Fieberg, & Matthiopoulos, 2012; McDonald, 2013), but as being justified on the basis of the required estimation of an odd. Logistic regression is the most common and straightforward method to estimate odds, but other probability models such as probit or complementary-log-log (clog-log) models could also be used, re-expressing results as odds (Fox, 2015). This would make no sense if all predictors are categorical variables, because in that case results are perfectly identical between logit, probit or clog-log models. When continuous predictors are involved, having the choice of models could offer more flexibility. In practice, it could however be more straightforward to use the logit link with either transformed predictors, polynomials or splines to obtain the desired level of flexibility. Regression coefficients are also more easily interpreted when using a logit link, as they represent odd-ratios, and results from regression with logit or probit links are usually very similar. Note that estimating SR in a regression context as proposed here, even when only one categorical predictor is present, naturally solves the difficulty of estimating confidence intervals for SR (Aho & Bowyer, 2015).

Overall, the reformulation of SR supports the common use of exponential RSF, but importantly highlights that other link functions could also be appropriate (see also Lele & Keim (2006) for a similar claim, on a different basis). It is worth emphasizing once more that the estimated odd is not the odd of the probability of selection, and that results from logistic regression in the context of habitat selection analyses should not be interpreted as a log-odd-ratio of the probability of selection (Lele et al., 2013; McDonald, 2013). This issue would disappear if the estimation process is pursued up to the estimation of SR.

Third, the reformulation of SR demonstrates that, contrary to current practice, the intercept of RSF models, which allows calculating the associated RSF score w_r_ for the reference habitat r, should not be overlooked. First, w_r_ allows the SR of the reference habitat to be estimated (eq. 7), which can be of interest in itself. Second, it is crucial to recognize that the estimation of the intercept affects the estimation of other parameters of the RSF. In particular, it is clear from equations above that the intercept aggregates a biological and a sampling process (see also Fieberg et al., 2010 and McDonald, 2013). Eq. 7 shows that 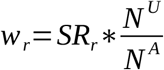, so that the value of the intercept will vary with the individual’s strength of selection for the reference habitat r, but also with the ratio of used/available locations in the dataset. This dependence of the intercept on a biological and a sampling process warrants some caution when fitting hierarchical (i.e., mixed) RSF models. For instance, it questions the recommendation made to use random intercepts to account for unbalanced sample size between individuals when fitting hierarchical RSF models (Gillies et al., 2006). Eq. 7 shows that what matters is actually not differences in the number of ‘used’ locations *N^U^* between individuals, but the ratio 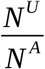. When this ratio is constant among individuals there is no need to account for differences in the number of ‘used’ locations. When this ratio varies between individuals, because for instance the same number of ‘available’ locations is chosen for all individuals whereas their number of ‘used’ locations vary, then random intercepts will account for both the variation in 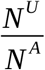 ratios *and* the difference in the strength of selection for the reference habitat (SR_r_). The fixed effect intercept will therefore be a meaningless average of these effects across individuals, and the parameter estimates for other habitats, which are estimated as difference from this reference habitat, will therefore also be biased. Similarly, when 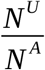 ratios vary between individuals, the random intercepts and their variance will not accurately quantify interindividual differences in the strength of habitat selection between individuals. In this case they should therefore not be used to assess ‘personalities’ for instance. This is especially true as, for categorical habitat variables, the variation in the strength of selection between individuals is spread between random intercepts (for the reference habitat) and random slopes (for other habitats). Fitting random intercepts and slopes is therefore required when fitting hierarchical RSF models (see also Muff et al., 2018 for a similar suggestion, based on a different ground). From personal experience, fitting such models could prove challenging (e.g., large computation time, frequent convergence issues) when categorical habitat variables are defined by many classes, or involved in a number of interactions with other variables. Two-stage approaches where one model is fit per individual and individual results later aggregated accounting for uncertainty in estimates may sometimes be easier to implement (Murtaugh, 2007; Fieberg et al., 2010).

Finally, the relevance of the intercept of RSF analyses also leads to questioning the current use of step-selection-function (SSF) analyses. SSF analyses are now routinely used when location data are collected at temporal resolution such that subsequent locations cannot be considered independent (Thurfjell, Ciuti, & Boyce, 2014). In SSF analyses, each used step is matched to a number of available steps, and SSF models are then fitted using conditional logistic regression with strata identifying the association between used and matched available steps (Fortin et al., 2005; Thurfjell et al., 2014). Because of the conditioning, conditional logistic regression models do not estimate an intercept (Duchesne, Fortin, & Courbin, 2010). This possibly explains why SSF models do not allow simulating utilization distribution when habitat selection is strong (Signer, Fieberg, & Avgar, 2017). Further work trying to bridge the gap between RSF and SSF models, or on other approaches for autocorrelated location data (Michelot et al., 2018), is urgently needed.

### Beyond the strength of selection, estimating the contribution of selection to habitat use

The SR is a natural and intuitive measure of the strength of habitat selection. It is well known however that the strength of habitat selection does not in itself reflects to what extent the amount of *use* of a habitat is modified by habitat selection (Lele et al., 2013). For instance, a highly selected habitat, as assessed by its SR, is used only moderately more than under random use if it is little available. This can be shown formally by considering that the selection effect (SE) on the use of habitat i can be estimated by:

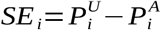

From eq. 1,

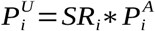

and therefore:

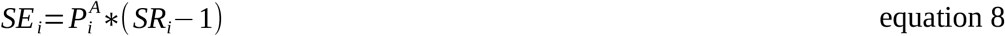

Eq. 8 makes clear that the selection effect of the amount of use of one habitat depends on both the strength of selection for this habitat and its availability. Once *SR_i_* have been estimated as suggested above, it is thus straightforward to estimate *SE_i_*, as 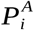 can be directly estimated using density estimation (this might require binning the continuous variables). Although *SE_i_* could also be expressed as a function of 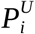 and *SR_i_*, its estimation would likely be less accurate as the estimation of 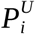 by density estimation is likely to be less precise that the one of 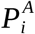, due to sample size. Note that, whereas SR is a unitless metric, SE is on the scale of use and availability measures. For instance, if one assumes that the number of locations in habitat i estimates the time spent in habitat i, then SEi estimates how much time spent in habitat i has been added or lost by selection compared to expectations under random use of habitats.

It is interesting to note that the SE index had already been proposed by Strauss (1979) as an index of selection (Lechowicz, 1982), and that an index equaled to the square of SE, termed marginality, is the basis of some statistical approaches of habitat selection (Ecological-Niche Factor Analysis: Hirzel et al. 2002; K-select analysis: Calenge, Dufour, & Maillard, 2005; General niche-environment system factor analysis: Calenge & Basille, 2008). More recently, myself and colleagues have emphasized that SE can be understood as the contribution of selection in determining use, and it provides a different, but complementary way, to measure habitat selection, compared to SR (Padié et al., 2015). SR estimates the effort made by the animal to select and use specific habitats, whereas SE estimates to what extent this effort actually modifies its habitat use, relatively to a random use of the available landscape. Thus, SR and SE do not estimate the same aspects of the habitat selection process and caution should be applied when comparing studies based on SR or RSF approaches with studies based on SE or marginality. Also, SE should be preferred when studying to what extent habitat selection affect demographic performances, as fitness is influenced by use, rather than selection itself (Gaillard et al., 2010).

Overall, the study of SR and SE will provide a comprehensive understanding of the strength of habitat *selection* and of its consequences for habitat *use.*

### An example: plains zebra habitat selection

To demonstrate the value of the approach described above, I conducted an analysis of dry-season habitat selection analyses of plains zebras *(Equus quagga),* using data from 19 individuals collected in Hwange National Park (Zimbabwe). I estimated the strength of habitat selection and the contribution of habitat selection on habitat use, using SR and SE respectively, in one of the management block of the Park. Habitats were defined using information on vegetation types (i.e., grassland, bushland or woodland) and distance to water. See Supplementary Appendix S1 for more information on the study area and the data.

I fitted an exponential RSF, which uses a logit link function, as this is the approach most commonly used. Coefficients from the RSF (Supplementary Appendix S1) showed that the strength of habitat selection increased with proximity to water, and was higher in grasslands that in the two other vegetation types. However, from such a RSF model, one could not know when, along the distance to water gradient, the strength of selection shifted from selection to avoidance. This information could however be critical in a management context, for example when deciding how to spread artificial waterholes in a landscape. Computing SR from the RSF model, following eq. 7, solved this issue and revealed that the shift from selection to avoidance occurred at ~ 2 km from water in bushland and woodland vegetation types, but at ~ 5 km from water in grasslands (Fig. 1A). Estimating the contribution of selection to habitat use provided further insights: the strong selection for habitats near water increased zebras’ use of grasslands (Fig. 1B), but had only a minor effect on the use of bushlands and woodlands because these were rare near water (Fig. 1B). The zebras’ habitat selection process affected the use of bushlands and woodlands only at intermediate (~ 3 to 7 km) distance to water, where these types of vegetation were less used than expected by chance. The general avoidance of areas located far from water also did not influence habitat use, compared to a random use, because these areas were rare in the landscape (Fig. 1B).

**Figure 1.**
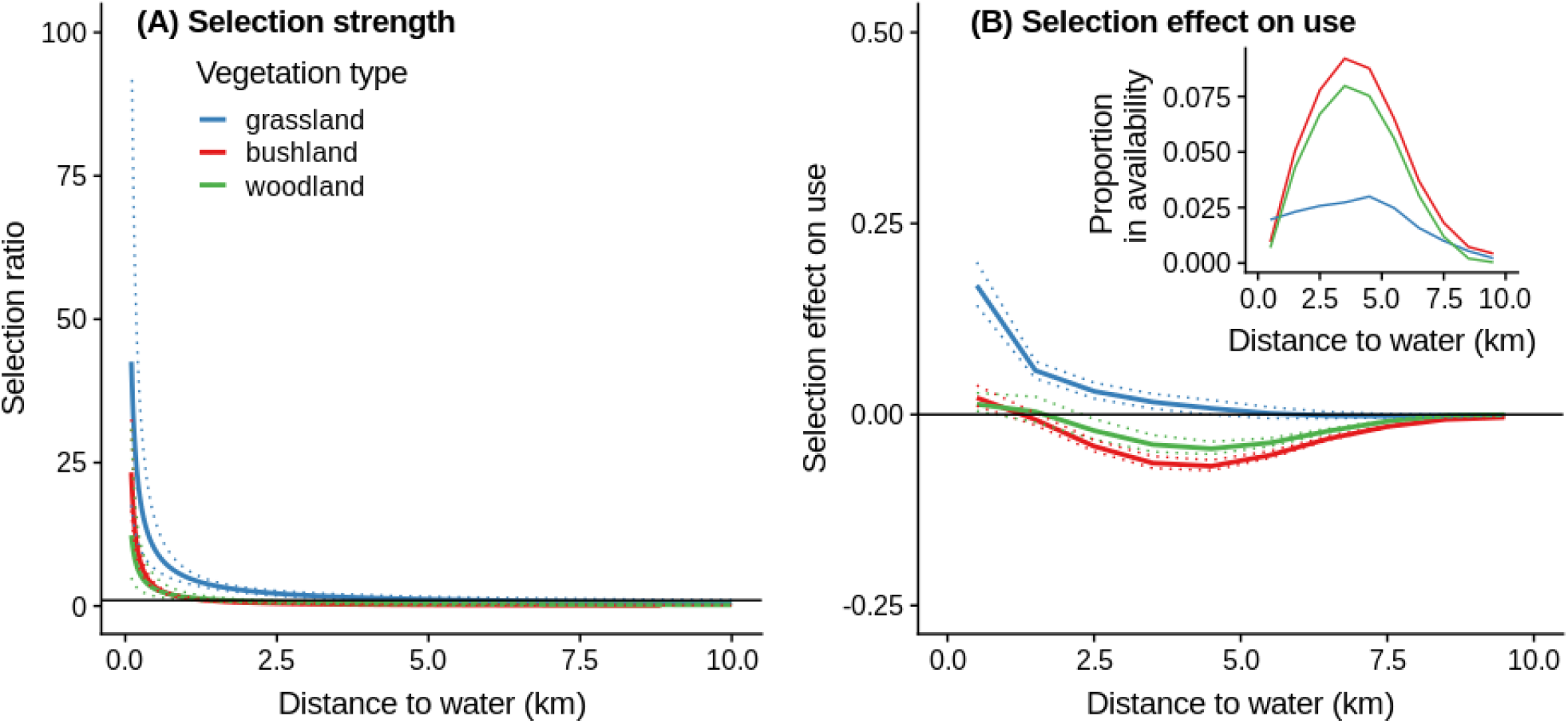
(A) Strength of zebra habitat selection, estimated using the selection ratio (SR), as a function of vegetation types and distance to water. Dotted lines show the 95% confidence intervals. (B) Contribution of habitat selection to habitat use, estimated using the selection effect (SE) metric (see text for details). Dotted lines show the 95% confidence intervals. The inset shows the distribution of habitats in the available landscape, estimated from data binned over 1 km bins.

## Conclusion

The reformulation of the SR presented here reveals that SR can be estimated in a regression context, even with continuous predictors, whereas it was initially the need to incorporate these predictors that led to the formulation of RSF (Mcdonald et al., 1990; Lele, 2009). It also clarifies some longstanding debates about RSF and data-selection/fitting practices. Finally, it shows that, with a trivially simple extra-step (i.e., multiplication of RSF scores by 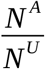), the interpretation of habitat selection analyses can be greatly clarified. Finally, it highlights the inter-relation between various metrics used to study habitat selection (i.e., SR, other selection indices, RSF scores, marginality). I thereby propose that SR and SE can be the unifying metrics of habitat selection, as together they offer a comprehensive view on the strength of habitat selection and its effect on habitat use.

## Acknowledgments

I thank C. Calenge, N. Courbin and S. Benhamou for constructive feedbacks on earlier versions of the manuscript. I thank R. Hedley for pointing out an error, made in previous versions of the manuscript, about the offset to be used in logistic regression. This work has been partly supported by the grant ANR-16-CE02-0001-01 of the French “Agence Nationale de la Recherche”.

## Supplementary Appendix

### S1. Presentation of the data used to analyze plains zebras habitat selection

The study was conducted using GPS data from zebra collared in Hwange National Park, Zimbabwe. The park vegetation is typical of dystrophic savannas: the vegetation is mostly made of bushland, interspersed by patches of woodlands and grasslands that are particularly common near waterholes (Chamaillé-Jammes, Fritz, & Madzikanda, 2009). For this study I used the 30-m resolution vegetation map produced by Arraut et al. (2018). I reduced the number of vegetation classes to three by pooling grasslands and bushed grasslands into a ‘grassland’ class, bushlands on Kalahari sands and scrublands on basalts into a ‘bushland’ class and deciduous woodland on Kalahari sands, deciduous woodland on basalts and evergreen woodlands into a ‘woodland’ class. The location of all permanent waterholes that retain water during the dry season is known (Chamaillé-Jammes, Fritz, & Murindagomo, 2007). I used GPS tracking data collected during the dry season (August to October) on 19 adult females living in different harems. Locations were originally acquired every 30 min or every hour but I reduced autocorrelation in the data by retaining only one location per day, the one closest to noon. See for instance Courbin et al. (2016, 2019) for further information about the zebra tracking study.

### S2. Statistical models and results

I studied zebra habitat selection by fitting resource selection function (RSF) models to estimate selection ratios (SR) and selection effect on use (SE). The RSF models were fitted in the use/available framework: I used the actual GPS locations as ‘used’ zebra locations, and drew ‘available’ locations randomly within the study area defined as the Main Camp management block of Hwange National Park (see Chamaillé-Jammes, Valeix, & Fritz 2007 for map). The ratio of available/used locations was 1000 for all individuals.

For the example provided here I fitted an exponential RSF model, which uses a logit link function, as it is the most commonly used RSF functional form. I used vegetation type (categorical variable with 3 categories, and ‘bushland’ as reference category), distance to water (continuous, log-transformed as we previously showed before that it greatly improved the fit of RSF models on these data (Courbin et al., 2016)) and the interaction between these two variables as predictors in the RSF model. I included zebra identity as random effects on the intercept and slopes for all predictors. All analyses were conducted with the R software v. 3.5.2 (R Core Team 2018) and the lme4 package v. 1.1.17 (Bates et al. 2015).

The exponential RSF model showed that the strength of habitat selection increased with proximity to water, and was higher in grasslands that in the two other vegetation types (Table S1).

**Table S1.** Estimates of fixed effects coefficients of the exponential resource selection function model. The reference vegetation type is ‘bushland’. P-values are based on the Wald Z-statistics and are reported as guidelines for interpretation only.

| Parameter | Estimate ± SE | P-value |
| --- | --- | --- |
| Intercept | -6.556 ± 0.124 | <0.001 |
| Vegetation type ‘grassland’ | 1.277 ± 0.142 | <0.001 |
| Vegetation type ‘woodland’ | 0.096 ± 0.198 | 0.629 |
| log(Distance to water) | -1.216 ± 0.154 | <0.001 |
| Vegetation type ‘grassland’ : log(Distance to water) | 0.292 ± 0.114 | 0.011 |
| Vegetation type ‘woodland’ : log(Distance to water) | 0.317 ± 0.125 | 0.011 |

However, from such a RSF model, one could not know when, along the distance to water gradient, the strength of selection shifted from selection to avoidance. Converting RSF scores to SR values solved this issue (see main text).

